# Nuclear m^6^A reader Ythdc1 regulates the scaffold function of LINE1 in mouse ESCs

**DOI:** 10.1101/2021.01.10.426065

**Authors:** Chuan Chen, Wenqiang Liu, Jiayin Guo, Yuanyuan Liu, Xuelian Liu, Jun Liu, Xiaoyang Dou, Rongrong Le, Yixin Huang, Chong Li, Lingyue Yang, Xiaochen Kou, Yanhong Zhao, You Wu, Jiayu Chen, Hong Wang, Bin Shen, Yawei Gao, Shaorong Gao

## Abstract

N^6^-methyladenosine (m^6^A) on chromosome-associated regulatory RNAs (carRNAs), including repeat RNAs, play important roles in tuning the chromatin state and transcription^1^. Among diverse RNA-chromatin interacting modes, the nuclear RNA scaffold is considered important for trans-interactions^2,3^ but has not yet been connected with m^6^A yet. Here, we found that Ythdc1 played indispensable roles in the embryonic stem cell (ESC) self-renewal and differentiation potency, and these roles highly depended on its m^6^A-binding ability. *Ythdc1* deficiency in ESCs resulted in decreased rRNA synthesis and the activation of 2-cell (2C) embryo-specific transcriptional program, and these observations recapitulated the transcriptome defects induced by dysfunction of the long interspersed nuclear element-1 (LINE1)-scaffold, which were unrelated to the direct targeting of Ythdc1. A detailed analysis revealed that Ythdc1 recognized m^6^A on LINE1 and was physically involved in the formation of the LINE1-Nucleolin partnership and the chromatin recruitment of Kap1. In summary, our study reveals a new link between m^6^A and the RNA scaffold and thus provides a new regulatory model for the crosstalk between RNA and the chromatin epigenome.

## Introduction

In eukaryotes, N^6^-methyladenosine (m^6^A) is the most prevalent internal modification of messenger RNAs (mRNAs) and long non-coding RNAs (lncRNAs)^4,5^, and this mark is involved in the regulation of various RNA-related processes, including pre-mRNA splicing, nuclear RNA transport, mRNA translation and RNA decay^6-10^. As an epigenetic mark, the reversible m^6^A modification of RNAs participates in the regulation of many essential biological events, including zygotic genome activation, cell fate transition and heat shock response^11-15^. An interesting issue which requires further investigations is whether m^6^A plays a role in RNA-chromatin interactions. To date, the diverse functions of chromatin-associated RNAs (caRNAs), including the functions of both coding and non-coding RNAs in gene regulation, have been revealed in different systems. In addition to cis-acting RNAs from nascent RNAs, a common trans-acting mode of RNA-chromatin interactions is through nuclear RNA scaffold; in this system, some RNA act as a scaffold to bind RNA-binding proteins (RBPs), and at least one component of this complex directly contacts with DNA-binding factors^2^. Recently, it was reported that RNA transcribed from long interspersed nuclear element-1 (LINE1) forms a LINE1-scaffold complex with Nucleolin (Ncl) and Kap1, which regulates the exit from the 2C-like state in mouse ESCs^3^, and these findings provide new insights for investigating RNA-chromatin interactions.

In a recent study, we and collaborators found the existence of m^6^A in chromatin-associated regulatory RNAs (carRNAs), including LINE1 RNAs. The m^6^A methylation on carRNAs mediated by Mettl3 could be further targeted by Ythdc1 for RNA decay and alterations in the chromatin state. However, quite different from Mettl3, we found that Ythdc1 plays essential roles in the maintenance of ESCs via a m^6^A-dependent mechanism; in addition, some Mettl3-independent m^6^A marks on LINE1 RNA could be defined, and these sites are also recognized by Ythdc1. Notably, the derepression of 2C-specific transcripts caused by LINE1-scaffold dysfunction was also observed in *Ythdc1*-deficient ESCs, and Ythdc1 physically interacts with the LINE1-Ncl-Kap1 complex. Further analysis revealed that Ythdc1 facilitates Kap1 recruitment to targets of the LINE1 scaffold, including 2C-specific retrotransposons, and this promotion contribute to the ESC identity.

## Results

### Indispensable Roles of Ythdc1 in Mouse ESCs

To explore the functional mechanism of *Ythdc1* in mouse ESCs, we constructed a conditional knockout ESC line (*Ythdc1*^flox/flox^ with *CreERT2*, referred to as *Ythdc1* cKO), and the *Ythdc1*^flox/flox^ line (referred to as *Ythdc1* f/f) served as the control (see Methods and Fig. S1A). Two rescue cell lines expressing WT Ythdc1 (referred to as wtRes) or m^6^A-binding-site-mutated Ythdc1 (referred to as W378A)^1,16^ were also constructed to identify the roles of m^6^A (Fig. 1A-B). As shown, treatment with 4-hydroxytamoxifen (4-OHT) deleted endogenous Ythdc1 protein in *Ythdc1* cKO ESCs (Figure S1B).

**Figure 1.**
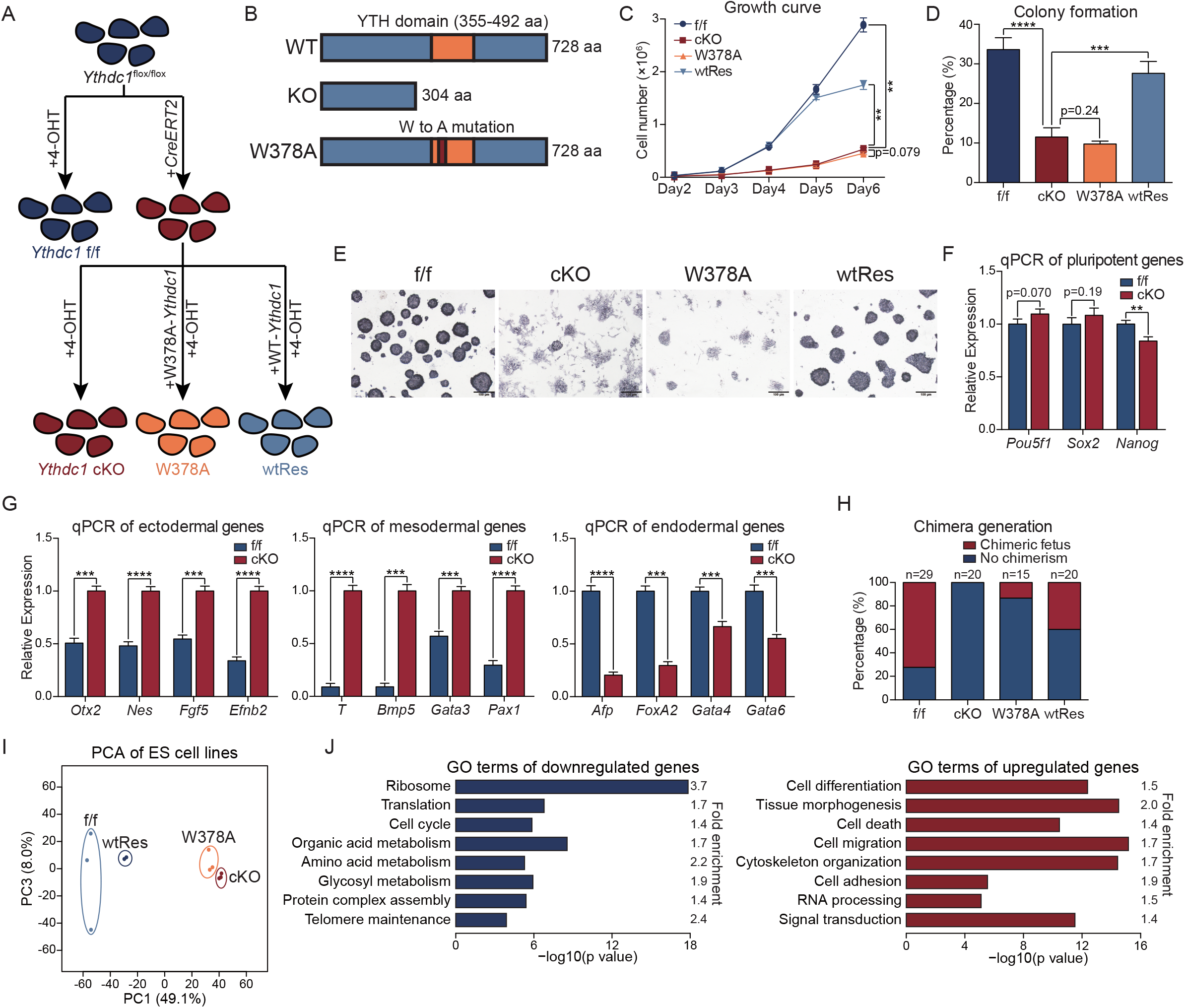
*Ythdc1* is essential for mouse ESCs. **(A)** Strategy for functional studies of Ythdc1 in ESCs. All cell lines were treated with 4-OHT for 3 days before harvesting to ensure the depletion of endogenous *Ythdc1*. **(B)** Schematic of wild-type (WT) Ythdc1, truncated Ythdc1 after the recombination (KO Ythdc1) and mutated Ythdc1 (W378A Ythdc1). **(C)** Growth curve showing that *Ythdc1* cKO and W378A ESCs exhibited a poor proliferation rate. Cell numbers on the last day were used to assess the significance. **(D and E)**. Colony formation ability of *Ythdc1* cKO and W378A ESCs was impaired, as revealed by AP staining. **(F)** qPCR analysis showing the relative RNA levels of key pluripotent markers in *Ythdc1* f/f and cKO ESCs. **(G)** qPCR analysis showing that EBs derived from *Ythdc1* cKO ESCs exhibited abnormal expression levels of differentiation markers 7 days after *in vitro* differentiation. **(H)** *Ythdc1* cKO and W378A ESCs exhibited a weak ability to generate chimeric mice. **(I)** Principal component analysis showing the transcriptome differences between each ESC line. **(J)** GO analysis of genes dysregulated in both *Ythdc1* cKO and W378A ESCs defined in Fig. S2D. Data are presented as means with SDs (n = 3 in C,F and G and n = 4 in D). Significance was calculated with unpaired two-tailed Student’s t test (** p < 0.01, *** p < 0.001, **** p < 0.0001).

Consistent with the unsuccessful establishment of homozygous *Ythdc1*-depleted ESCs^17^, we found that *Ythdc1*-deficient ESCs could hardly support cell maintenance, as demonstrated by a markedly reduced proliferation rate and impaired colony formation ability (Fig. 1C-E and S1C). An EdU incorporation assay showed that *Ythdc1* cKO ESCs failed to enter the S phase (Fig. S1D-E), and the apoptosis rate of *Ythdc1* f/f and cKO cells was comparable (Fig. S1F). However, despite *Ythdc1* depletion, key pluripotent markers appeared to be normal, with the exception of slight decreases in the *Nanog* RNA and protein levels (Fig. 1F and S1G). Notably, mutant Ythdc1 could hardly rescue these maintenance defects. We then evaluated the developmental potency of *Ythdc1* cKO ESCs through *in vitro* and *in vivo* differentiation. As shown, embryoid bodies (EBs) derived from *Ythdc1* cKO ESCs were markedly fewer and smaller (Fig. S1H-I), and the markers of all three germ layers were expressed abnormally in *Ythdc1* cKO EBs (Fig. 1G). We introduced transgenic red fluorescence protein (RFP) into the four ESC lines and checked their integrating abilities in chimeric mice. Strikingly, no chimera was observed with the use of *Ythdc1* cKO ESCs, and W378A ESCs could hardly contribute to chimeric embryos either (Fig. 1H and S1J-K). These results suggest that Ythdc1 is essential for the self-renewal and proper differentiation of ES cells, and these roles are highly related to its m^6^A-recognition ability.

We then performed RNA-seq analysis of each line, and found that the transcriptomes of *Ythdc1* cKO and W378A ESCs shared more similarity, which was quite different from those of *Ythdc1* f/f or wtRes ESCs (Fig. 1I and S2A). Furthermore, the differentially expressed genes (DEGs) in *Ythdc1* cKO compared with f/f ESCs (546 upregulated and 1373 downregulated genes, Fig. S2B and Table S1) can be largely corrected by WT Ythdc1 but not the W378A mutant protein (Fig. S2C). We then identified the transcriptome defects shared by *Ythdc1* cKO and W378A ESCs (Fig. S2D and Table S1) and found that the downregulated genes are closely related to ribosome- and translation-associated processes and that the upregulated genes are involved in cell differentiation and cell adhesion (Fig. 1J and S2E). This finding led us to hypothesize that the regulation of Ythdc1 in ESC is dependent on its m^6^A-recognition ability. Notably, the large-scale repression of ribosomal genes might lead to reduced rRNA synthesis, as was also observed in LINE1-knockdown (KD) ESCs, and is considered responsible for the observed growth defects^3^.

### Ythdc1 Targets LINE1 RNA

We subsequently questioned whether the transcriptional impact was related to the direct binding of the Ythdc1 protein. Through nuclear RNA-binding protein immunoprecipitation and sequencing (RIP-seq), we identified 12237 peaks of Ythdc1, and Ythdc1 distribution was enriched near the center of m^6^A peaks (Fig. S3A). We found that Ythdc1 binding was highly enriched in intronic regions and endogenous retrotransposons (Fig. 2A and S3B), which was consistent with the published regulatory roles of Ythdc1 on alternative splicing and repeat RNAs^1,9^. For coding genes, we found that most genes with Ythdc1 RIP peaks in exons also possessed peaks in introns (Fig. S3C), and a decreased usage of exons adjacent to Ythdc1 RIP peaks (Fig. 2B) and a slightly reduced expression level of target genes was detected after *Ythdc1* depletion (Fig. 2C). However, these target genes were not enriched in ribosome- or translation-associated terms, and most of these genes were not included in the main transcriptome defects shared by *Ythdc1* cKO and W378A ESCs (Fig. S3D-E). These results indicate that the dysregulated transcriptome resulting from *Ythdc1* deficiency is not directly related to the cis-regulation of Ythdc1-targeted RNA.

**Figure 2.**
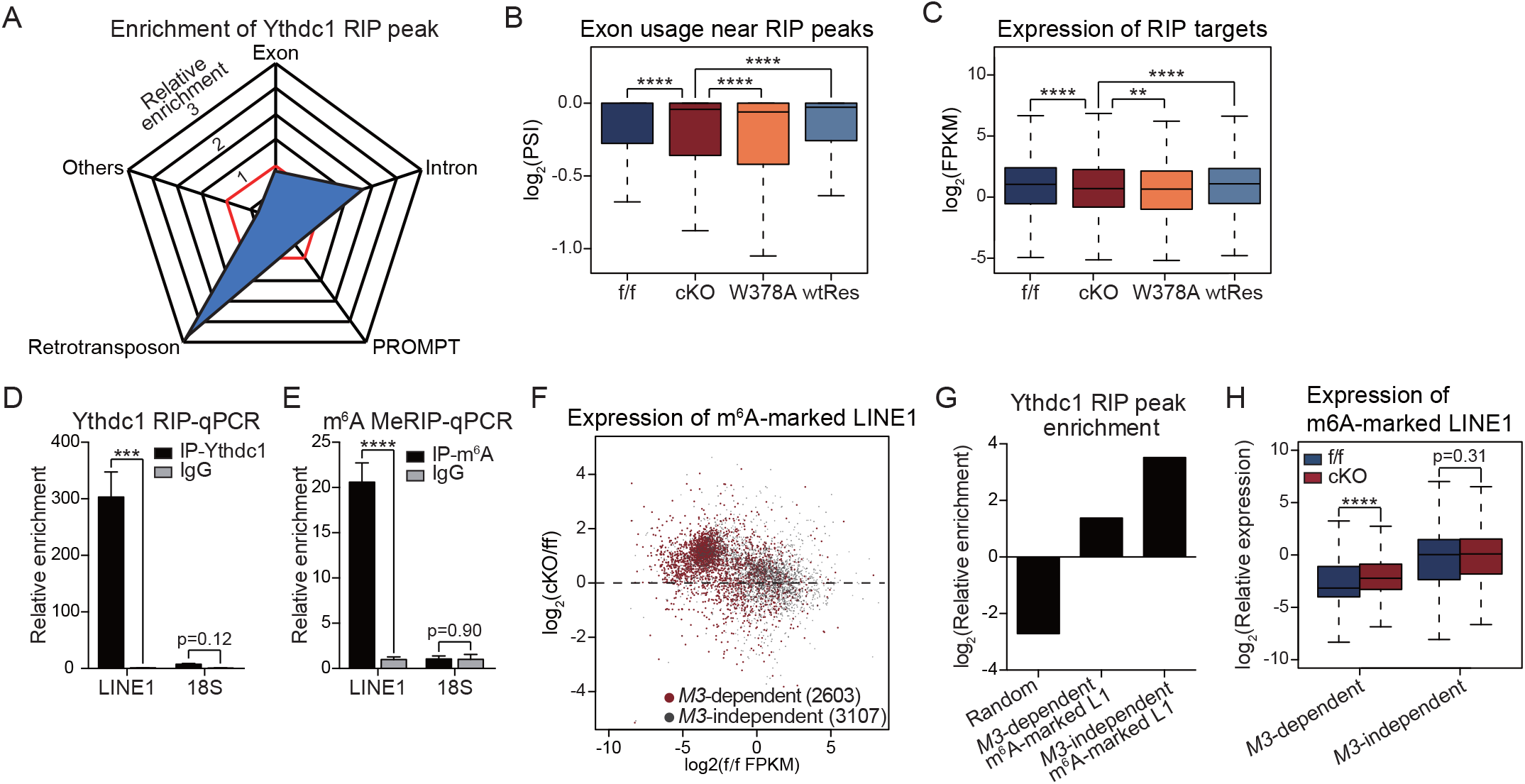
Targets of Ythdc1 in mouse ESCs. **(A)** Ythdc1 RIP peaks preferred to localize at introns and retrotransposons in genome. **(B)** Boxplot showing that the usage of exons (calculated as PSI, see Methods) adjacent to Ythdc1 RIP peaks was decreased upon *Ythdc1* depletion. **(C)** Boxplot showing the expression levels of Ythdc1 RIP targets in each ESC line. A total of 941 coding genes possessing RIP peaks (fold enrichment > 5) in their exons or introns were defined as RIP targets. **(D and E)** Ythdc1 RIP-qPCR and methylated RNA IP-qPCR showing the preference of Ythdc1 binding and m^6^A deposition on LINE1 transcripts in ESCs, respectively. Relative enrichment was calculated as the percent of input relative to the negative control antibody IgG. Two independent reactions for each IP were perfromed. Data are presented as means with SDs (n = 3 technical replicates). **(F)** Scatter plot showing the expression levels of m^6^A-marked LINE1 caRNAs in *Ythdc1* f/f ESCs (x-axis) and the expression fold change upon*Ythdc1* depletion (y-axis). Expression levels of individual copies were quantified as FPKM using Cufflinks. **(G)** Ythdc1 RIP peaks were enriched on both Mettl3-dependent and Mettl3-independent m^6^A-marked LINE1 transcripts in ESCs. Mettl3-dependent m^6^A peaks were detected in WT ESCs but were not found in *Mettl3* KO cells, and Mettl3-independent m^6^A peaks were retained despite *Mettl3* KO. M3, *Mettl3* gene. L1, LINE1 copies. **(H)** Boxplot showing that Mettl3-dependent m^6^A-marked LINE1 caRNAs were upregulated and that Mettl3-independent m^6^A-marked LINE1 caRNAs remained unchanged in *Ythdc1* cKO ESCs. Significance (*** p < 0.001, **** p < 0.0001) was calculated with two-tailed Student’s t test (paired in B,C and H and unpaired in E and F).

We then investigated the binding of Ythdc1 on endogenous retrotransposons. We calculated the relative enrichment of Ythdc1 RIP peaks and m^6^A peaks^1^ on major classes of retrotransposons. As shown, LINE1 possesses highly enriched Ythdc1 RIP peaks and m^6^A peaks (Fig. S3F), and these findings were further confirmed by RIP-qPCR and methylated RNA immunoprecipitation (MeRIP)-qPCR, respectively (Fig. 2D-E). In an earlier study, we found that Mettl3 deposits m^6^A modifications on carRNAs, including LINE1, and facilitates RNA decay with Ythdc1 and the NEXT complex^1^. However, the loss of *Mettl3* did not lead to a substantial loss in proliferation ability and did not prevent ESCs from forming compacted colonies, which was different from depletion of *Ythdc1* (Fig. S3G-3I). These findings indicate that Ythdc1 might regulate some RNAs in a Mettl3-independent manner. We then defined the LINE1 copies on chromatin with Mettl3-dependent and Mettl3-independent m^6^A peaks (Fig. 2F) based on the published caRNA m^6^A-seq data of *Mettl3* KO and wild-type (WT) ESCs^1^. Interestingly, although both groups of LINE1 transcripts were highly enriched with Ythdc1 binding (Fig. 2G), they presented different characteristics upon *Ythdc1* depletion. The RNA level of LINE1 copies on chromatin with Mettl3-dependent m^6^A peaks is relatively low in the control and was significantly increased in *Ythdc1* cKO ESCs; in contrast, LINE1 caRNAs with Mettl3-independent m^6^A peaks were abundant in the control, and their RNA level was not impacted by the depletion of *Ythdc1* (Fig. 2F and 2H). These results suggest that Ythdc1 can bind LINE1 RNA and play new regulatory roles in addition to facilitating the decay of caRNAs.

### Ythdc1 Depletion Resembles LINE1-Scaffold Dysfunction

In mouse ESCs, LINE1 RNA acts as a nuclear scaffold for recruiting Ncl/Kap1, repressing the 2C program, activating rRNA synthesis and promoting ESC self-renewal^3^. Considering the similar ribosome-associated and maintenance defects of *Ythdc1* cKO ESCs, we wondered whether the binding of Ythdc1 was related to the scaffold function of LINE1. We first defined DEGs in LINE1 ASO ESCs using published RNA-seq data^3^ and found that over 50% of these DEGs showed similar dysregulation upon *Ythdc1* depletion (Fig. 3A and S4A). A recent study identified LINE1-targeted genes and LINE1 sequence-enriched genes through chromatin isolation by RNA purification followed by sequencing (ChIRP-seq), and these genes are repressed by the LINE1-scaffold complex^18^. We found that *Ythdc1* deficiency in ESCs also led to similar upregulation of both LINE1-targeted and LINE1-enriched genes (Fig. S4B), which indicated that Ythdc1 might be important for the repression function of LINE1.

**Figure 3.**
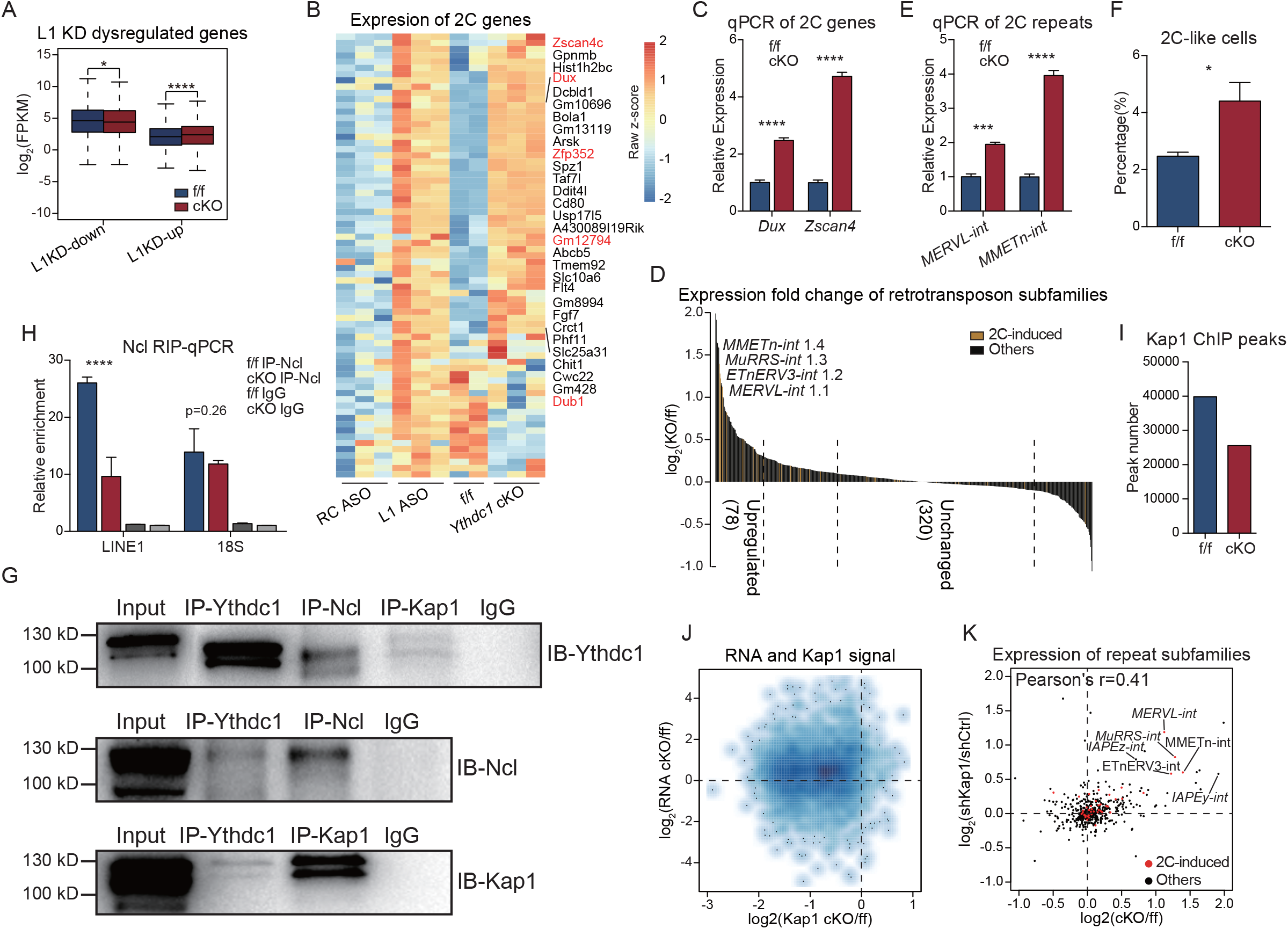
Ythdc1 is involved in the functions of the LINE1-scaffold complex. **(A)** Boxplot showing that DEGs detected in LINE1 ASO ESCs were similarly dysregulated in *Ythdc1* cKO ESCs. **(B)** Heatmap showing that many 2C-induced genes were upregulated in both LINE1 KD ESCs and *Ythdc1* cKO ESCs. Genes with log_2_ (fold change) > 1 (2C-tdTomato^+^ versus 2C-tdTomato^-^) were considered 2C-induced genes. RC ASO, reverse complement of LINE1 ASO (served as a negative control). L1 ASO, LINE1 RNA KD by ASO. **(C and E)** qPCR analysis showing that representative 2C-induced genes and retrotransposons were upregulated in *Ythdc1* cKO ESCs. **(D)** Many 2C-induced retrotransposons were upregulated in *Ythdc1* cKO ESCs. A total of 611 detectable subfamilies were ranked in descending order according to their expression fold change (*Ythdc1* cKO versus f/f). Subfamilies with log_2_(fold change) > 0.3 were considered *Ythdc1* cKO-upregulated repeats (Table S2), and subfamilies with |log_2_(fold change)| < 0.1 were considered unchanged after *Ythdc1* depletion. 2C-induced repeats were identified as retrotransposons with log_2_(fold change) > 1 (2-cell versus oocytes), and representative subfamilies are labeled in the plot. **(F)** Percentage of 2C-like cells was increased after *Ythdc1* depletion in the ESC population. **(G)** Co-IP showing that endogenous Ythdc1 interacted with both Kap1 and Ncl proteins in mouse ESCs. **(H)** Ncl RIP-qPCR showing that the association of Ncl with LINE1 RNA was attenuated in *Ythdc1* cKO ESCs. Relative enrichment was calculated as the percent of input relative to the negative control antibody IgG. Two independent reactions for each IP were perfromed. **(I)** Kap1 ChIP peaks were reduced in *Ythdc1* cKO ESCs. **(J)** Kap1 ChIP signals (x-axis) were decreased and RNA levels (y-axis) were increased on Kap1 ChIP peaks overlapped with LINE1 RNA-targeted loci in *Ythdc1* cKO ESCs. **(K)** Many retrotransposons, including 2C-induced retrotransposons, were upregulated upon *Ythdc1* depletion (x-axis) or *Kap1* KD (y-axis) in ESCs. Pearson’s correlation coefficient was calculated in R. shCtrl, control shRNA expressed. shKap1, *Kap1* shRNA expressed. Data are presented as means with SDs in C,E,F and H (n = 3). Significance (* p < 0.05, *** p < 0.001, **** p < 0.0001) was calculated with two-tailed Student’s t test (unpaired in C,E and H and paired in F).

As reported, the LINE1-Ncl-Kap1 complex represses the 2C transcriptional program and promotes rRNA synthesis in mouse ESCs^3^. We then defined 2C-induced genes and retrotransposons using published data^3,19^ and observed consistent upregulation of 2C-induced genes in LINE1 ASO and *Ythdc1* cKO ESCs, including *Dux* and *Zscan4* (Fig. 3B-C and S4C). Moreover, most of the 2C-specific retrotransposons were also upregulated in *Ythdc1* cKO ESCs (Fig. 3D-E and Fig. S4D; Table S2), and this effect could not be rescued by W378A Ythdc1 (Fig. S4E). We next introduced a reporter of the 2C-like state, a *Zscan4* promoter-driven green fluorescence protein (GFP)^20^, into *Ythdc1* cKO ESCs. Consistent with the upregulation of the 2C-specific transcripts, the proportion of GFP-positive cells was significantly increased upon *Ythdc1* depletion (Fig. 3F and S4F), which indicated that the loss of Ythdc1 promotes the transition of ESCs into the 2C-like state. Moreover, *Ythdc1* depletion also led to an obvious decrease in rRNA levels, and this decrease could not be recovered in W378A ESCs (Fig. S4G). Taken together, these results indicate that *Ythdc1* deficiency recapitulates the dysregulation of the 2C transcription program and rRNA synthesis caused by LINE1 KD, which suggests that the scaffold function of LINE1 is restrained by the depletion of *Ythdc1*.

### Ythdc1 Regulates LINE1-Ncl and Recruits Kap1 to Chromatin

In addition to the direct binding of Ythdc1 to LINE1 RNA, we hypothesized that Ythdc1 might also interact with proteins in the LINE1-Ncl-Kap1 complex. We performed co-immunoprecipitation (Co-IP) with ESCs, and found that Ythdc1 interacted with both Ncl and Kap1 proteins (Fig. 3G), and the interaction between mouse Ythdc1 and Ncl proteins was also observed when these two proteins were exogenously expressed in HEK293T cell (Fig. S5A). In addition, Ncl might serve as a mediator to connect Kap1 and Ythdc1 as the interaction between the latter two proteins was more obvious when Ncl was also co-expressed in HEK293T cells (Fig. S5B-C).

Because the RNA or protein levels of *Ncl* and *Kap1* were not impacted by *Ythdc1* depletion (Fig. S5D-E) and few genomic sites were found to be directly targeted by the Ythdc1 protein (data not shown), we next questioned whether Ythdc1 could regulate the formation of the LINE1-Ncl-Kap1 complex. We performed RIP-qPCR and found that the enrichment of Ncl protein toward LINE1 RNA was significantly decreased in *Ythdc1* cKO ESCs, but its binding toward 18S rRNA was not impacted (Fig. 3H). We then performed chromatin immunoprecipitation followed by sequencing (ChIP-seq) for Kap1 and found that the peak number and relative ChIP signal were decreased in *Ythdc1* cKO ESCs (Fig. 3I and S5F). A detailed analysis revealed that many of the impacted Kap1 peaks were located in retrotransposons (Fig. S5G), and most of the Kap1 occupancy decreases on LINE1-targeted loci were combined with transcription upregulation (Fig. 3J), which suggested that the derepression of LINE1-targeted loci in *Ythdc1* cKO ESCs might result from insufficient recruitment of Kap1 to chromatin. To confirm this inference, we checked the transcriptome changes induced by *Kap1* KD and found that the LINE1-targeted and LINE1-enriched genes were also upregulated (Fig. S5H). Moreover, 2C genes and retrotransposons were derepressed in *Kap1* KD ESCs, resembling the defects observed in *Ythdc1* cKO ESCs (Fig. 3K and S5I-J). Taken together, our data indicate that when directly targeting the m^6^A on LINE1 RNA, Ythdc1 facilitates the formation of the LINE1-Ncl partnership and promotes the chromatin recruitment of Kap1 to silence LINE1-targeted 2C genes and retrotransposons.

## Discussion

In summary, our data supplement the working model in which Ythdc1 binds the m^6^A on LINE1 RNA and regulates the formation of the LINE1-Ncl-Kap1 complex to repress the 2C program, which ensures the appropriate transcriptome and developmental potency of ESCs. In contrast to the Ythdc1-mediated co-transcriptional direction of m^6^A toward H3K9me2 demethylation^26^, RNA modifications on the nuclear scaffold might be involved in complicated chromatin regulation on multifarious loci, which is highly efficient and powerful. Our findings further explore the cross-layer interaction model between RNA and chromatin and highlight the potential regulatory roles of RNA modifications on carRNAs in tuning transcription and the chromatin state in mouse ESCs.

## Methods

### Cell line Generation and Cell Culture

The ESC clone for *Ythdc1* cKO (C57BL/6N-*Ythdc1*^tm1a(EUCOMM)Wtsi^) was purchased and microinjected into mouse blastocysts. The resulting chimeric mice were crossed with flipper mice to excise the FRT flanked selection cassette. *Ythdc1*^flox/flox^ ESCs were derived from *Ythdc1*^flox/flox^ blastocysts. 2×10^5^ ESCs were transfected with 200 ng PB-CAG-CreERT2-P2A-puromycin and 100 ng PBase by electroporation using Lonza Nucleofector-4D. 24 hr later, electroporated cells were treated with 1 µg/ml puromycin to screen for Ythdc1^flox/flox^ ESC clones stably expressing *CreERT2*. To generate rescue ESC lines, *Ythdc1*^flox/flox^; *CreERT2* cells were electroporated with PB-CAG-Ythdc1-P2A-blasticidin and PB-CAG-Ythdc1-W378A-P2A-blasticidin, respectively. 24 hr after the electroporation, 1 µg/mL puromycin and 10 µg/mL blasticidin was added to generate stable *Ythdc1*^flox/flox^; *CreERT2*; WT-Ythdc1 and Ythdc1^flox/flox^; *CreERT2*; W378A-Ythdc1 ESC lines.

ESCs were cultured in ESC medium (DMEM-high glucose, 15% FBS, 1×Nucleosides, 1 mM L-Glutamine, 1×Non Essential Amino Acids, 0.1 mM 2-Mercaptoethanol, 1000 U/ml LIF) supplemented with 3 mM CHIR-99021 and 1 mM PD0325901. To knock out the endogenous *Ythdc1* in *Ythdc1*^flox/flox^; *CreERT2* and rescue lines, 2.5 µM 4-OHT was added into the culture medium which made the Cre recombinase enter the nuclei and trigger the deletion of exon 5-7 at *Ythdc1* locus. ESCs of each line were harvested for further investigations after 72-hr treatment of 4-OHT.

### Generation of Knockout Mouse Embryos and Isolation of ICM

To ensure the effective deletion of *Ythdc1* in most embryos, four single guide RNAs (sgRNAs; Table S3) targeting exon 4 of *Ythdc1* gene were designed. The sequence for each sgRNA was cloned into the sgRNA expression vector pUC57 and in vitro transcription was then performed using MEGAshortscript T7 Transcription Kit. The Cas9 expression construct pST1374-Cas9-N-NLS-Flag-linker was linearized and transcribed using mMESSAGE mMACHINE T7 ULTRA Transcription Kit. B6D2F1 female mice (approximately 8-week-old) were superovulated and mated with B6D2F1 male mice before zygotes were collected from the oviducts. The harvested zygotes were microinjected with a mixture containing 100 ng/µL Cas9 mRNA and 50 ng/µL each sgRNA, and the embryos were then cultured in G-1 PLUS medium. To test the knockout efficiency, we used 50 injected embryos as the PCR template and amplified genomic fragments spanning exon 4 of *Ythdc1* gene (primers are listed in Table S3). The PCR product was ligated to pClone007 vector and the resulting product was then transformed into E.Coli bacteria. We observed large fragment deletions in all of 30 sequenced bacterial colonies, indicating *Ythdc1* was efficiently knocked out by the CRISPR/Cas9 system in most embryos.

For the isolation of ICM cells, blastocysts were incubated in Ca^2+^-free Chatot-Ziomek-Bavister (CZB) medium for 30 min to disrupt the cell-cell junction. ICM cells were then distinguished and collected according to their sizes and shapes with the aid of a piezo-driven micromanipulator.

### Alkaline Phosphatase Staining

AP staining was performed using the BCIP/NBT Alkaline Phosphatase Color Development Kit. ESCs were fixed with 4% paraformaldehyde for 10 min at room temperature before they were washed with PBS twice and stained with the BCIP/NBT Solution following manufacturer’s protocol.

### Growth Curve and Colony Formation Assay

ESCs were dissociated by 0.05% trypsin-EDTA and the cell density was determined with a counting chamber. Cells were then plated in 12-well plates at 20000 cells per well. 2 days, 3 days, 4 days, 5 days and 6 days after the passaging, cell number in each well was counted to plot the growth curve.

To analyze the ability of forming colonies, ESCs were digested and counted before they were seeded into 24-well plates at 500 cells per well. 3 days later, cells were fixed and AP staining was performed as described above. AP^+^ colonies were then counted under an inverted microscope to calculate the percentages of cells which were able to form colonies in each line.

### Apoptosis Assay and EdU Incorporation

ESCs were digested and collected for the apoptosis assay, which was performed according to the manufacturer’s instructions of Annexin V-FITC Apoptosis Detection Kit. For the cell cycle analysis, 10 µM EdU was added to the culture medium 2 hr before the cells were harvested. The EdU labeling was then performed following the instructions of Click-iT Plus EdU Alexa Fluor 647 Flow Cytometry Assay Kit. The percentages of apoptotic cells and S-phage cells were determined by fluorescence-activated cell sorting (FACS) and the results were analyzed through CytExpert software.

### Embryoid Body Differentiation

ESCs were trypsinized and resuspended at 50000 cells/mL in LIF-free ESC medium. Hanging drops containing 20 µL of cell suspension were made on the lids of 10 cm-dishes and 10 mL of PBS was added in each dish below the drops. 2 days later, the resulting EBs were transferred to non-adherent 6-well plates and cultured in LIF-free medium for another 5 days before they were harvested for gene expression detection.

### Generation of Fluorescent Protein-Labeled ESC Lines and Chimeric Mice

The pSicoR-Ef1a-mCherry-Puro plasmid together with the lentiviral packaging plasmids psPAX2 and pMD2.G were introduced into HEK293T cells using VigoFect reagent, and the transfected cells were cultured for 48 hr to allow virus assembly and expansion. The supernatant medium containing viruses was mixed with 10% PEG 8000 and rotated for approximately 12 hr at 4 °C. The mixture was then pelleted and resuspended with ESC medium, and the concentrate was subjected to infect ESCs of each line. After the infection, the RFP^+^ cells were screened and collected using FACS.

To generate chimeric mice, the RFP-labeled ESCs were trypsinized and approximately 10 ESCs were microinjected into each E3.5 blastocyst derived from the ICR mouse strain. These embryos were then implanted into the uteri of female pseudo pregnant ICR mice. 11 days after the implantation, the surrogate mothers were dissected and fetuses were harvested for further investigations. The extent of chimerism in each fetus was determined by the percentage of RFP^+^ cells using FACS analysis on the skin tissues.

### Detection of the 2C-Like Cell Proportion within ESC Culture

The sequence encoding a *Zscan4c* promoter-driven GFP were constructed into the Fugw plasmid, which was then introduced into the cKO ESCs through lentivirus infection as described above. Cell lines with identical genetic materials were established from individual ES colonies after the infection, and three independent lines were used for subsequent analysis. To compare the 2C-like cell proportion before and after Ythdc1 depletion, the three lines were cultured in ESC medium without or with 4-OHT for 3 days and subjected to FACS analysis. Cells with credible GFP expression were considered to be in a 2C-like state.

### Short Hairpin RNA (shRNA)-Mediated Gene Silencing

The sequence encoding each shRNA was constructed into the shRNA expression vector pSicoR-RFP, which was then introduced into WT ESCs through lentivirus infection as described above. The successfully infected ESCs were screened by FACS. shRNA sequences are listed in Table S3.

### Western Blotting

Cells were harvested and subjected to 1×Omni-Easy Protein Sample Loading Buffer for protein denaturing. Proteins were then separated by 10% SDS/PAGE and western blotting was performed by standard procedures. Primary antibodies were used as follows: anti-Gapdh (1:3000), anti-Ythdc1 (1:1000), anti-Nanog (1:2000), anti-Kap1 (1:2000), anti-Ncl (1:2000), anti-H3 (1:3000), anti-H3K9me3 (1:3000). HRP-conjugated secondary antibodies (1:5000) were then applied. Protein signals were measured using High-sig ECL Western Blotting Substrate and visualized with ChemiDoc MP imaging system.

### Co-Immunoprecipitation

Approximately 5×10^6^ cells were pelleted and sheared in lysis buffer (10 mM HEPES, pH 7.4, 150 mM NaCl, 2 mM EDTA, 1% Nonidet P 40, 0.5 mM DTT, protease inhibitor). The cell lysate was then incubated with specific antibody-bound Dynabeads Protein A/G in IP buffer (50 mM HEPES, pH 7.4, 200 mM NaCl, 2 mM EDTA, 0.05% Nonidet P-40, 0.5 mM DTT, protease inhibitor) for 8 hr at 4 °C with rotation. Bound immunocomplexes were washed twice with IP buffer and 4 times with high-salt IP buffer (IP buffer with 500 mM NaCl) before they were eluted by heat in 1×Omni-Easy Protein Sample Loading Buffer and subjected to immunoblotting.

### Reverse Transcription and Quantitative Real Time-PCR

ESCs were disrupted in TRIzol Reagent and total RNAs were isolated by chloroform extraction coupled with isopropanol precipitation. RNAs of ICM cells were also isolated using TRIzol Reagent and chloroform, with the exception that 1/10 volume of 3 M NaAc and 1 µL glycogen was added to the aqueous phase of each sample. RNAs were then precipitated by isopropanol and washed twice with 75% ethanol before they were eluted with nuclease-free water. cDNA was then synthesized using All-In-One RT MasterMix. qPCR was carried out using TB Green Premix Ex Taq II and monitored by 7500 Fast Real-Time PCR System, and three technical replicates were performed for each sample. Relative expression level of each gene was normalized to the reference gene *Hprt* for ESC samples or *H2afz* for embryo samples. qPCR primers for tested genes are listed in Table S3.

### Ribosomal RNA-Free Total RNA-Seq

Total RNAs were extracted as described above. For ESCs, 1 µg RNA per sample was subjected to rRNA elimination and RNA-seq library generation using VAHTS Total RNA-seq (H/M/R) Library Prep Kit following manufacturer’s instructions. For ICM cells, purified RNAs were subjected to library generation using SMARTer Stranded Total RNA-Seq Kit. Libraries were sequenced on the Illumina NovaSeq 6000 platform with paired ends and 150-bp read lengths at Berry Genomics.

### Nuclear RNA-Binding Protein Immunoprecipitation

The RIP for Ythdc1 protein was performed according to manufacturer’s instructions of Nuclear RNA-Binding Protein Immunoprecipitation Kit. Briefly, 5×10^6^ WT ESCs were cross-linked in 0.3% formaldehyde for 10 min at room temperature and neutralized by 1×glycine. The fixed cells were pelleted by centrifugation and lysed with nuclei isolation buffer. The isolated nuclei were then pelleted and subjected to sonication using the Covaris M220 with conditions as follows: peak power 75, duty factor 20, cycles/burst 200, 5 min. 1/20 of the sonication product was saved as the input and the rest was incubated with Ythdc1 or Ncl antibody-coated Dynabeads Protein A/G for 12 hr with rotation at 4 °C. After the incubation, beads were washed and the immunoprecipitated complex was eluted as instructed. RNAs of the saved input and the IP product were extracted and subjected to either library generation using VAHTS Total RNA-seq (H/M/R) Library Prep Kit, or reverse transcription for qPCR. The paired-end sequencing was performed as described above. Primers for RIP-qPCR are listed in Table S3.

### Chromatin Immunoprecipitation

For ESCs, 5×10^6^ cells per sample were fixed in 1% formaldehyde and the cross-link was stopped by 125 mM glycine. The crosslinked cells were pelleted and washed twice with PBS before they were subjected to lysis buffer (50 mM Tris-HCl, pH 8.0, 10 mM EDTA, 1% SDS, protease inhibitor). The sonication was then performed with the following conditions: peak power 75, duty factor 20, cycles 200, 10 min. The product was cleared by centrifugation, and 1/100 of the supernatant was saved as the input. Antibodies for Kap1 or H3K9me3 were pre-bound to Dynabeads Protein A/G, and the sheared chromatin was incubated with the beads in RIPA buffer (10 mM Tris-HCl, pH 7.5, 140 mM NaCl, 1% Triton X-100, 0.1% SDS, 0.5 mM EGTA, 1 mM EDTA, protease inhibitor) for 12 hr at 4 °C with rotation. After the immunoprecipitation, beads were washed twice with RIPA buffer, 4 times with high-salt RIPA buffer (RIPA buffer with 500 mM NaCl) and once with TE buffer (10 mM Tris-HCl, pH 8.0, 10 mM EDTA). The immune complexes were then eluted in elution buffer (20 mM Tris-HCl, pH 7.5, 50 mM NaCl, 1% SDS, 5 mM EDTA, 0.2 mg/mL protease K) at 68 °C with shaking for 2 hr. For DNA purification, the input and IP samples were mixed with an equal volume of DNA Extraction Reagent, and 1/10 volume of 3 M NaAc together with 1 µL glycogen was added to the aqueous phase of each sample before DNAs were precipitated by isopropanol. Sequencing libraries from the purified DNAs were generated using HyperPlus Library Preparation Kit. Primers for ChIP-qPCR are listed in Table S3.

The ultra-low-input native ChIP-seq was performed as previously described^27,28^ to capture the H3K9me3 status in ICM cells. Briefly, the harvested ICM cells were subjected to nuclei extraction buffer (10 mM Tris-HCl, pH 8.0, 140 mM NaCl, 5 mM MgCl2, 0.6% Nonidet P-40, 1 mM PMSF, protease inhibitor). The MNase Master mix (0.6 U/µL MNase, 1×MNase master buffer, 2 mM DTT, 5% PEG 6000) was then added and the mixture was incubated at 25 °C for 10 min to allow MNase digestion. The reaction was stopped by 10 mM EDTA, and 0.1% Triton X-100 together with 0.1% DOC was added to lyse the nuclear membrane. The released chromatin was then diluted with ChIP buffer (10 mM Tris-HCl, pH 8.0, 90 mM NaCl, 2 mM EDTA, 0.1% Triton X-100, 0.1% DOC, 0.1 mM PMSF, protease inhibitor). After 1/20 of the reaction was saved as the input, the rest was incubated with H3K9me3 antibody-coated Dynabeads Protein A/G for 12 hr at 4 °C with rotation. The IP samples were washed twice with low-salt wash buffer (20 mM Tris-HCl, pH 8.0, 150 mM NaCl, 2 mM EDTA, 1% Triton X-100, 0.1% SDS, protease inhibitor) and twice with high-salt wash buffer (wash buffer with 500 mM NaCl). The washed beads were then incubated in hot elution buffer (100 mM NaHCO_3_, 1% SDS) for 2 hr at 65 °C with shaking. The eluted DNAs were further purified and subjected to sequencing library generation as described above. Paired-end 150-bp sequencing was also performed on ChIP libraries.

### RNA-Seq Data Processing and Analysis

Sequencing reads were trimmed by Trim Galore (v 0.6.4, with parameters: --phred33, --illumina, --clip_R1 9, --clip_R2 9, --paired) to remove adapter sequences before they were aligned to Mouse genome version mm9 by the unique mapping of Tophat2 (v 2.1.1)^29^, which reported the alignment with the best alignment score if multiple alignments were mapped for a given read.Expression level of individual genes in each sample was quantified by fragments per kilobase of exon model per million reads mapped (FPKM) using Cufflinks (v 2.2.1)^30^, and genes with low expression levels (averaged FPKM < 0.1) were ignored. Differentially expressed genes were defined as genes with fold change > 2 using the cuffdiff function of Cufflinks. Gene ontology (GO) analysis for functional annotation of gene sets was performed on the DAVID website, which provided fold enrichment of each term and p values showing the significance, and clusters were summarized to several representative terms. Usage of individual exons in each sample was calculated as percent spliced in index (PSI), which was the ratio between reads including or excluding the designated exons^31^. Gene set enrichment analysis (GSEA) was performed in the software (v 4.0.1)^32^, which provided normalized enrichment scores and nominal p-values. To quantify RNA level for different kinds of retrotransposons, genomic regions of each repeat subfamily were defined; the aligned files were normalized according to the sequencing depth; the relative expression level of a subfamily was calculated based on the number of reads falling on its copies using the intersect function of BEDTools (v 2.20.1)^33^. To compare the read coverage at specific sites between samples, the normalized BAM files were converted to BedGraph files using the genomecov function of BEDTools, which were further converted to BigWig files using bedGraphToBigWig^34^ and visualized in Integrative Genomics Viewer (IGV)^35^. The normalized BigWig files were also used to quantify RNA signal on designated regions with bigWigSummary,

### RIP-Seq and ChIP-Seq Data Processing and Analysis

Raw reads from RIP-seq libraries were trimmed and uniquely mapped to the reference genome mm9 using Tophat2 as described above. The output BAM files were used to define RIP peaks by the callpeak function of MACS2 (v 2.1.1)^36^ with the following parameters: -g mm, -f BAM, --BDG, --QVALUE 0.05, --SPMR. The intersected peaks defined in both of the replicates were considered as high confident peaks. To compare the enrichment of RIP peaks on different elements, genomic regions of promoter upstream transcripts (PROMPTs, 2 kb upstream of the transcription start site), exons, introns and retrotransposons were defined; the percentage of peaks falling on each element was then calculated, which is further divided by the percentage of input reads falling on the corresponding element, and this ratio is regarded as the relative enrichment for the element. The relative enrichment of RIP and m^6^A peaks^1^ on individual repeat subfamilies was calculated based on similar methods.

ChIP-seq reads were trimmed and uniquely mapped to mm9 using Bowtie2 (v 2.2.9)^37^ with the default parameters. The output SAM files were converted to sorted BAM files using SAMtools (v 1.3)^38^, and duplicate reads were removed using the MarkDuplicates tool of Picard (v 2.22.2). The filtered files were further normalized according to the sequencing depth. Peak calling was performed using MACS2 and the relative peak enrichment was computed as described above. The BedGraph files generated by MACS2 were further converted to BigWig files using bedGraphToBigWig (v 4)^34^. Signal tsracks were also visualized in IGV. The normalized BigWig files of ChIP and input samples were used to quantify the averaged signal at designated regions by bigWigSummary, and the ratio (IP/input) was regarded as ChIP signal intensity.

### Statistical analysis

All statistical analyses were performed in R (v 3.5.1). Numbers of replicates in each test are labeled in the graph or stated in the legend, and the error is reported in the graph as the standard deviation (SD). The statistical significance (* p < 0.05, ** p < 0.01, *** p < 0.001, **** p < 0.0001) between groups was determined by two-tailed Student’s t test. For sequencing data, average values of replicates are presented.

---

**Figure S1.**
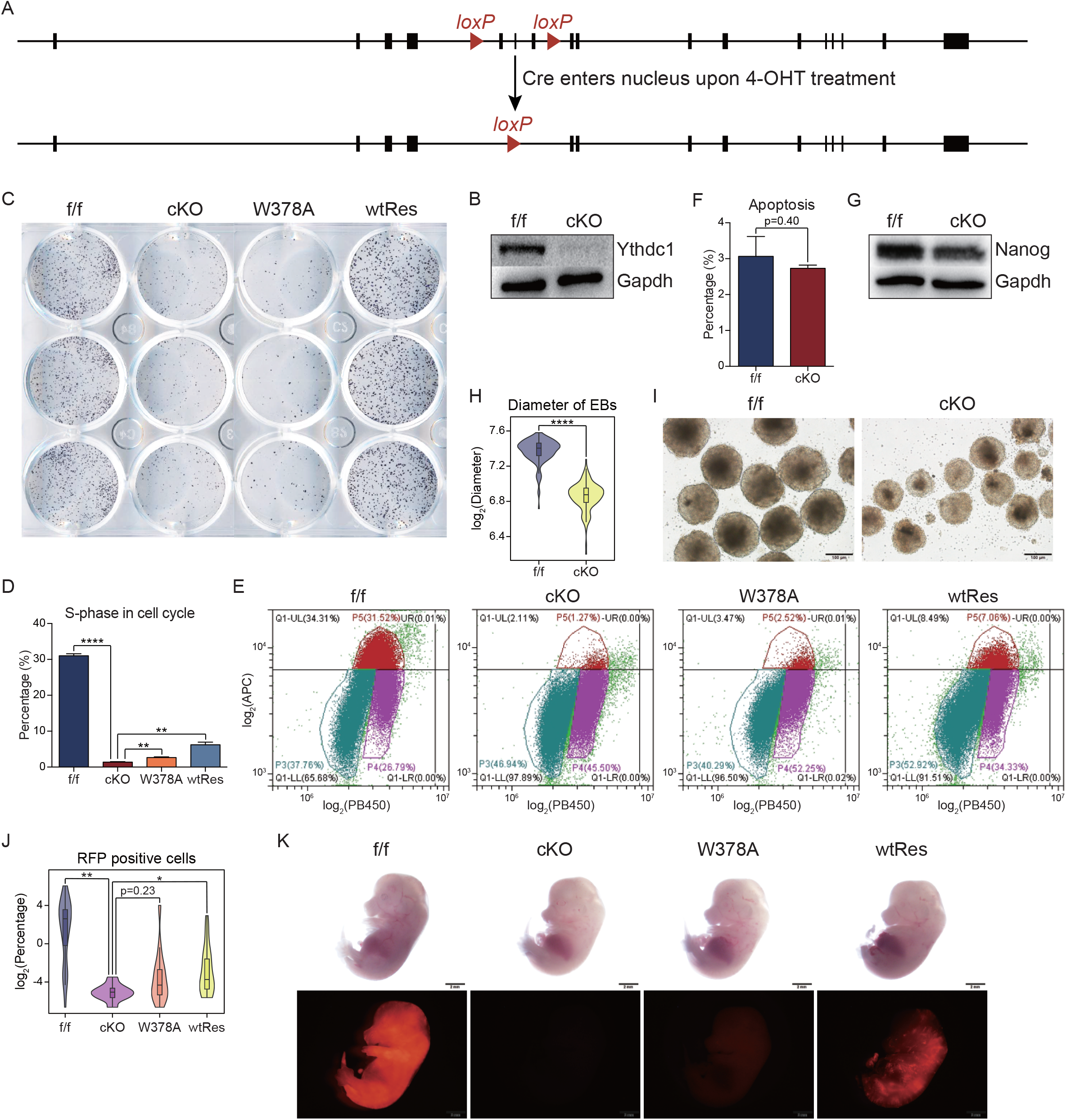
Growth and differentiation defects caused by *Ythdc1* deficiency in ESCs, related to Fig. 1. **(A)** Strategy for *Ythdc1* conditional knockout in ESCs. Upon 4-OHT treatment, Cre recombinase enters the nuclei and triggers the deletion of exon 5-7 at *Ythdc1* locus in the cKO ESCs. **(B)** Western blotting showing that most of Ythdc1 protein was depleted in *Ythdc1* cKO ESCs. **(C)** *Ythdc1* cKO and W378A ESCs exhibited impaired colony formation ability (related to Fig. 1D-E). **(D and E)** *Ythdc1* cKO and W378A ESCs could hardly enter the S phase. DNA content (PB450, x-axis) and EdU incorporation (APC, y-axis) are shown in **E**. **(F)** Percentage of early apoptotic cells (defined as Annexin V^+^ cells) was comparable in *Ythdc1* f/f and cKO ESCs. **(G)** Western blotting showing Nanog protein was slightly reduced in *Ythdc1* cKO ESCs. **(H and I)** Diameters of embryoid bodies formed by *Ythdc1* cKO ESCs were smaller than those formed by f/f ESCs 4 days after *in vitro* differentiation. Violin plot showing diameters of 132 EBs for each line in **H**. **(J and K)** *Ythdc1* cKO and W378A ESCs could hardly contribute to chimeric embryos (related to Fig. 1H). Violin plot showing percentage of RFP^+^ cells in E14.5 fetus derived from the injected blastocysts. Data are presented as means with SDs in D and F (n = 3). Significance was calculated with unpaired two-tailed Student’s t test (* p < 0.05, ** p < 0.01, **** p < 0.0001).

**Figure S2.**
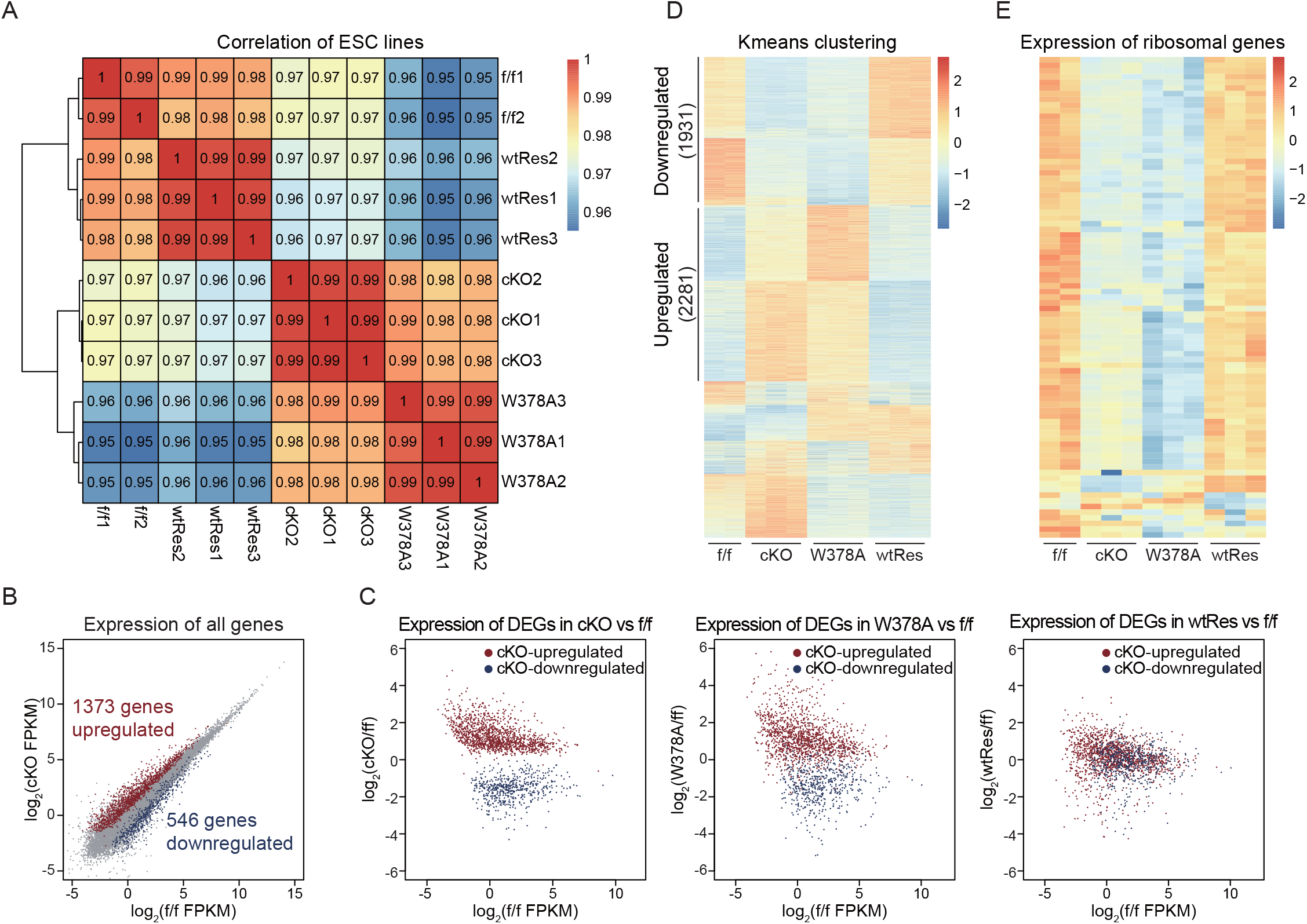
*Ythdc1* deficiency leads to transcriptome defects in ESCs, related to Fig. 2. **(A)** Hierarchical clustering showing that transcriptomes of *Ythdc1* cKO and W378A ESCs were similar, which were relatively different from those of f/f and wtRes ESCs. Pearson’s correlation between each replicate is labeled in the plot. **(B)** Scatter plot showing the expression level of all expressed genes in *Ythdc1* f/f (x-axis) and cKO (y-axis) ESCs. Differentially expressed genes in *Ythdc1* cKO ESCs (Table S1) are labeled in the plot. **(C)** Expression level of DEGs defined in *Ythdc1* cKO ESCs (Table S1) could be corrected by WT Ythdc1 protein but not the W378A mutant protein. **(D)** K-means clustering (k = 8; Table S1) identified genes simultaneously dysregulated in *Ythdc1* cKO and W378A ESCs. **(E)** Heatmap showing that ribosomal genes were extensively downregulated in *Ythdc1* cKO and W378A ESCs. A total of 83 Rpl and Rps genes are included in the plot.

**Figure S3.**
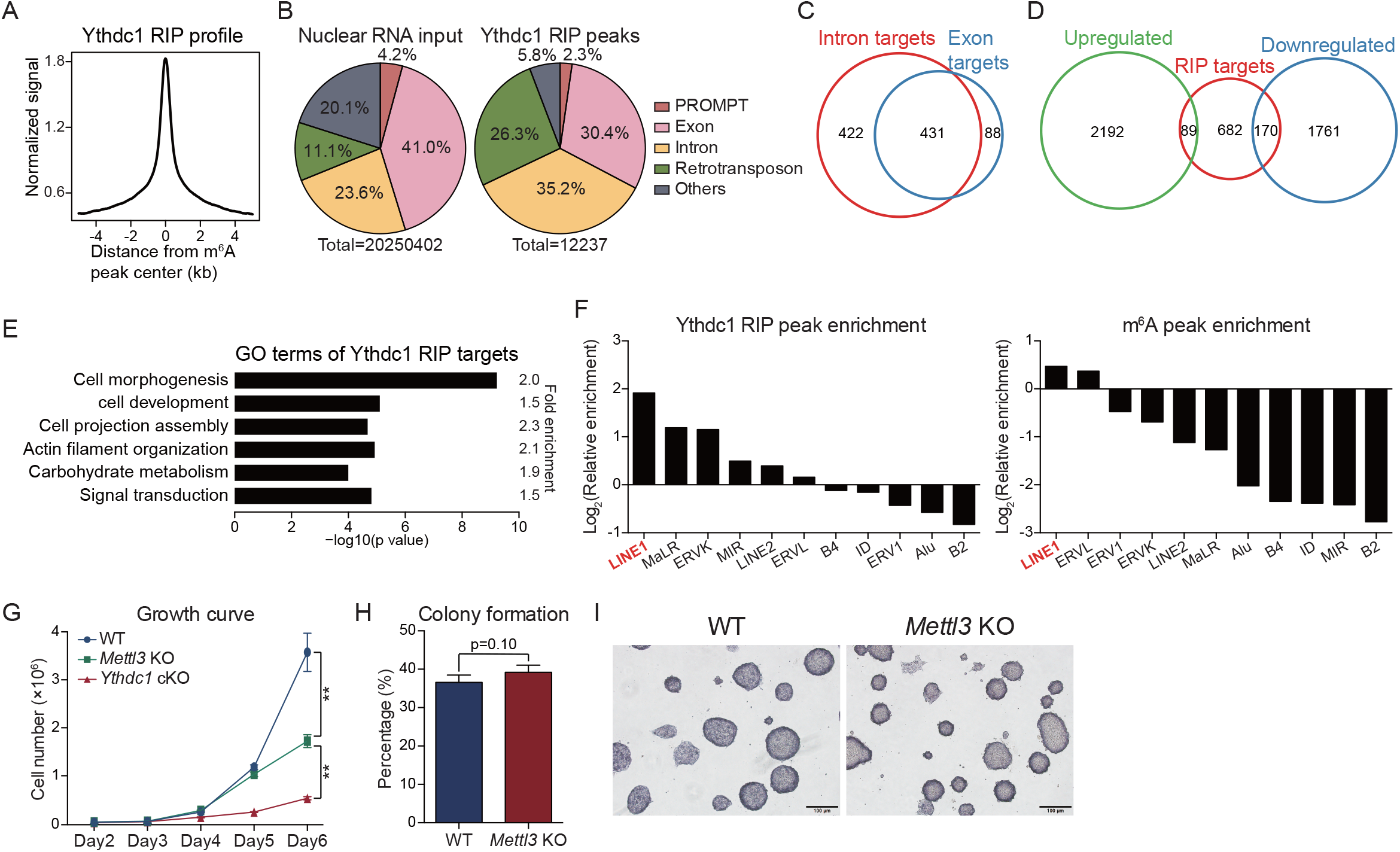
Targets of Ythdc1 protein revealed by RIP-seq analysis, related to Fig. 2. **(A)** Distribution of Ythdc1 RIP-seq signal around the center of m^6^A peaks. **(B)** Pie chart showing the distribution of nuclear RNA input reads (left) and Ythdc1 RIP peaks (right) on different genomic elements in ESCs. **(C)** Venn diagram showing the overlap of genes targeted by Ythdc1 at exonic regions (exon targets) and genes targeted by Ythdc1 at intronic regions (intron targets) in ESCs. **(D)** Venn diagram showing the overlap of Ythdc1 RIP targets and dysregulated genes defined in Fig. S2D. **(E)** GO analysis of Ythdc1 RIP targets. **(F)** Both Ythdc1 RIP peaks (left) and m^6^A peaks (right) were enriched on transcripts of LINE1. Representative subfamilies of retrotransposons are included in the graph. **(G)** Growth curve showing that the self-renewal ability was more severely impaired in *Ythdc1* cKO ESCs compared to *Mettl3* KO ESCs. Cell numbers on the last day were used to assess the significance. **(H)** Colony formation ability of WT and *Mettl3* KO ESCs was comparable. **(I)** Morphology of colonies was not impacted by *Mettl3* depeletion in ESCs, as revealed by AP staining. Data are presented as means with SDs (n = 3 in G and n = 4 in H). Significance was calculated with unpaired two-tailed Student’s t test (** p < 0.01).

**Figure S4.**
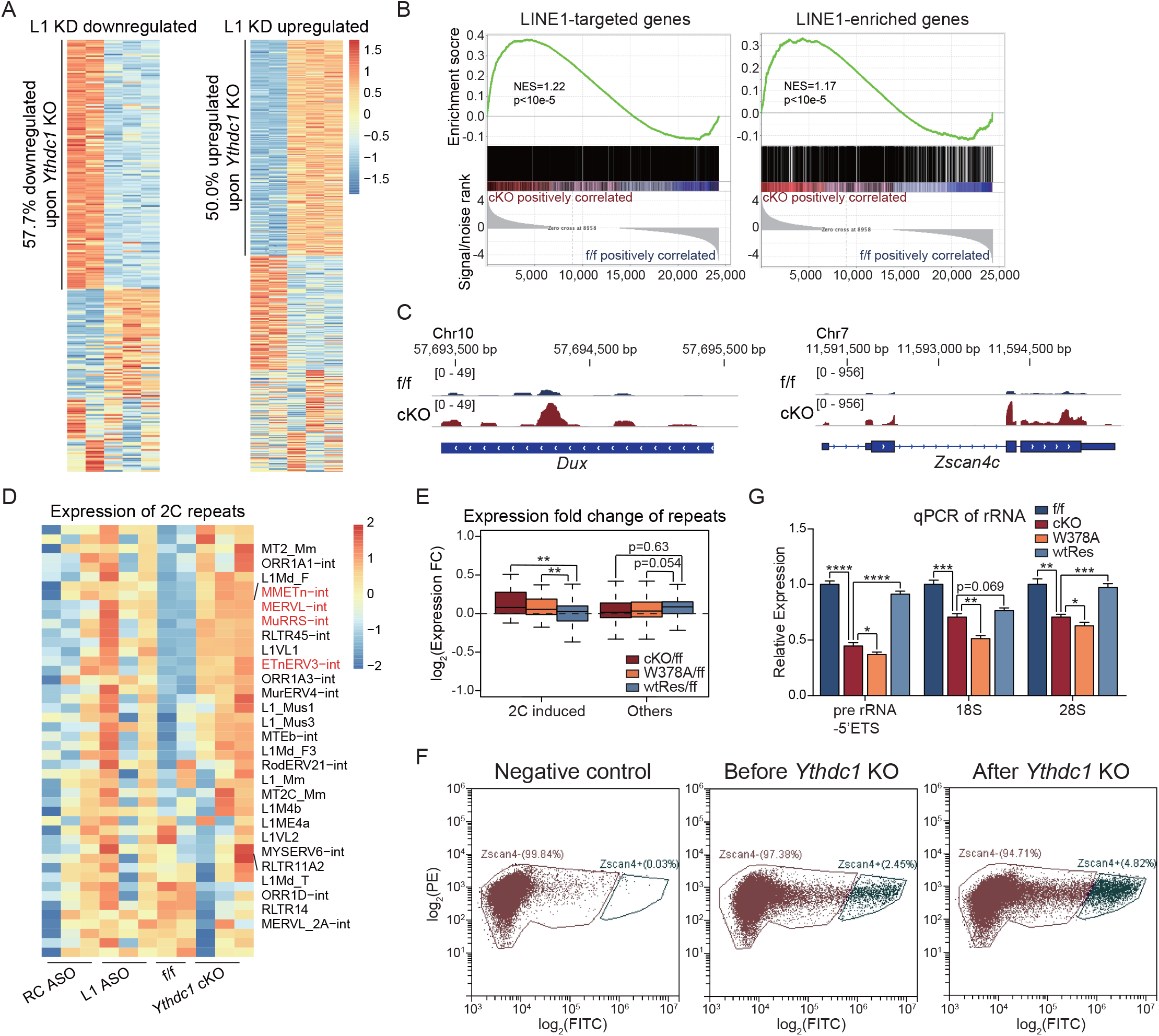
*Ythdc1* deficient ESCs share transcriptome defects with LINE1 knockdown ESCs, related to Fig. 3. **(A)** Heatmap showing that many DEGs defined in LINE1 ASO ESCs (Percharde et al., 2018) were similarly dysregulated in *Ythdc1* cKO ESCs. **(B)** GSEA showing the global upregulation of LINE1 RNA-targeted genes (left) and LINE1 sequence-enriched genes (right) in *Ythdc1* cKO ESCs. These genes were defined by Lu et al. (2020). NES, normalized enrichment score. **(C)** IGV tracks showing that the RNA-seq read coverage at *Dux* (left) and *Zscan4c* (right) loci was higher in *Ythdc1* cKO ESCs. **(D)** Heatmap showing many 2C-induced retrotransposons (Macfarlan et al., 2012) were upregulated in both LINE1 ASO and *Ythdc1* cKO ESCs. **(E)** Boxplot showing 2C-induced retrotransposons were upregulated in *Ythdc1* cKO and W378A ESCs, and this effect could be rescued by WT Ythdc1 protein. FC, fold change. **(F)** Fluorescence signal plot showing that the percentage of 2C-like cells was increased after *Ythdc1* depletion in a representative cKO line. The 2C-like cell population was determined according to the GFP signal distribution (x-axis) in a negative control line without the *Zscan4c* promoter-GFP transgene. **(G)** qPCR analysis showing that rRNA level was decreased in *Ythdc1* cKO and W378A ESCs. Data are presented as means with SDs (n = 3).

**Figure S5.**
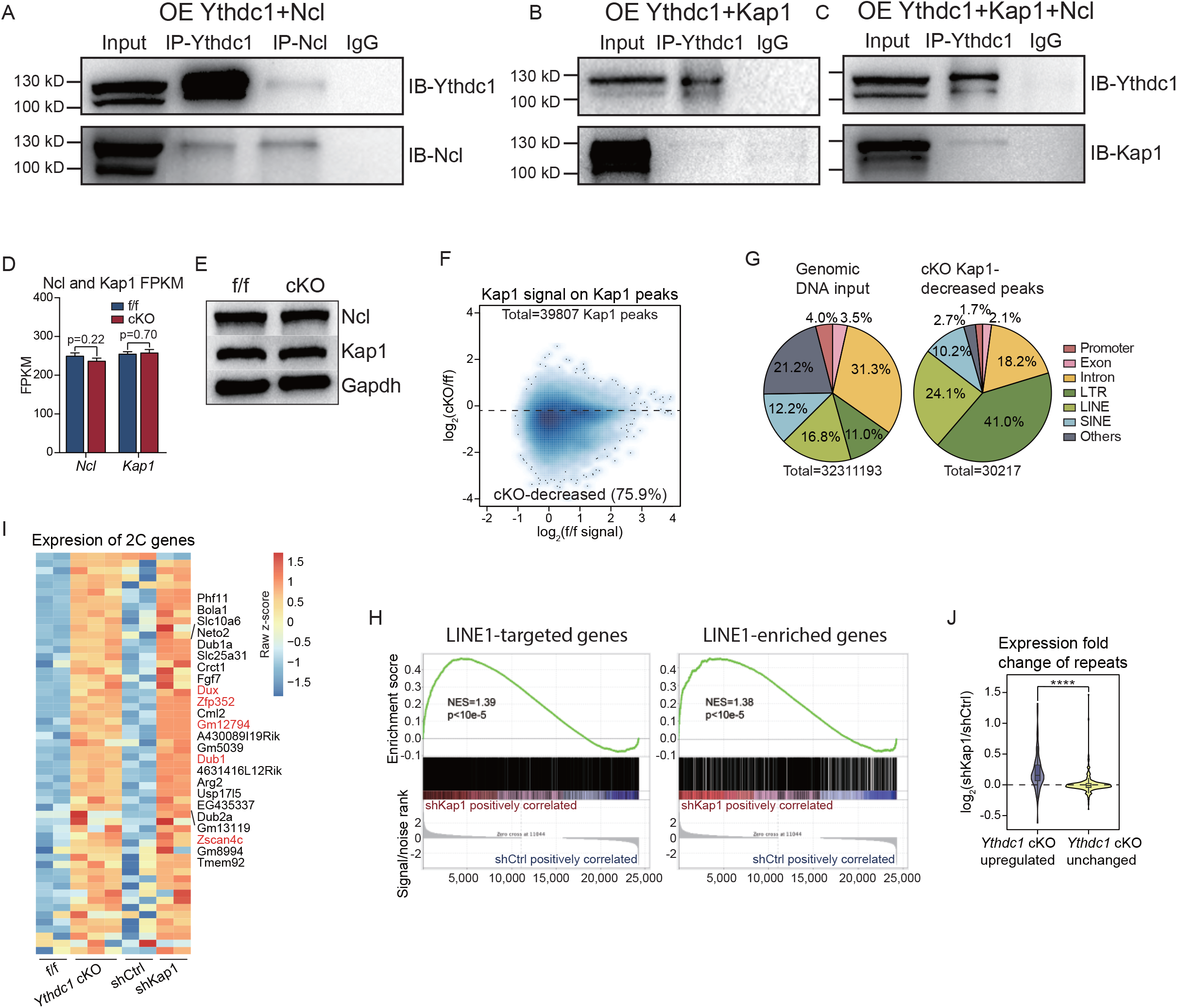
Roles of Kap1 is restrained upon *Ythdc1* depletion in ESCs, related to Fig. 3. **(A)** Co-IP showing the interaction between exogenously expressed mouse Ythdc1 and Ncl proteins in HEK293T cells. **(B and C)** Co-IP showing the association of exogenously expressed mouse Ythdc1 and Kap1 proteins was more obvious when mouse Ncl protein was co-expressed in HEK293T cells. **(D and E)** RNA and protein level of *Ncl* and *Kap1* was comparable in *Ythdc1* f/f and cKO ESCs. Data are presented as means with SDs in **D** (n = 3). **(F)** Scatter plot showing the Kap1 ChIP signals in *Ythdc1* f/f ESCs (x-axis) and the signal fold change upon *Ythdc1* depletion (y-axis) on Kap1 ChIP peaks. cKO-decreased Kap1 peaks were defined as Kap1 peaks with log_2_(fold change) < -0.2. **(G)** Pie chart showing the distribution of genomic DNA input reads (left) and cKO-decreased Kap1 peaks defined in **F** (right) on different genomic elements in ESCs. LTR, long terminal repeat. SINE, short interspersed nuclear element. **(H)** GSEA showing the global upregulation of LINE1 RNA-targeted genes and LINE1 sequence-enriched genes (Lu et al., 2020) upon *Kap1* KD in ESCs. NES, normalized enrichment score. **(I)** showing that many 2C-induced genes (Percharde et al., 2018) were consistently upregulated in *Ythdc1* cKO ESCs and *Kap1* KD ESCs. **(J)** Violin plot showing that the expression levels of *Ythdc1* cKO-upregulated retrotransposons (defined in Fig. 3D) were also increased upon *Kap1* KD in ESCs.

